# SpatialSort: A Bayesian Model for Clustering and Cell Population Annotation of Spatial Proteomics Data

**DOI:** 10.1101/2022.07.27.499974

**Authors:** Eric Lee, Kevin Chern, Michael Nissen, Xuehai Wang, IMAXT Consortium, Chris Huang, Anita K. Gandhi, Alexandre Bouchard-Côté, Andrew P. Weng, Andrew Roth

## Abstract

Emerging spatial proteomics technologies have created new opportunities to move beyond quantifying the composition of cell types in tissue and begin probing spatial structure. However, current methods for analysing such data are designed for non-spatial data and ignore spatial information. We present SpatialSort, a spatially aware Bayesian clustering approach that allows for the incorporation of prior biological knowledge. SpatialSort clusters cells by accounting for affinities of cells of different types to neighbours in space. Additionally, by incorporating prior information about cell types, SpatialSort outperforms current methods and can perform automated annotation of clusters.

## Background

Recently developed high throughput spatial protein expression profiling technologies can perform highly multiplexed phenotyping of single cells, while preserving the spatial organization of tissues. Examples of such technologies include imaging mass cytometry (IMC)^1^, multiplexed ion beam imaging (MIBI)^2^, and co-detection by indexing imaging (CODEX)^3^. These technologies have the capacity to quantify dozens of protein markers at single cell resolution in-situ. This provides an opportunity to enhance studies of cellular heterogeneity, by going beyond the quantification of cellular composition and allowing for direct inference of cell to cell interactions from spatial context.

A key step when analysing spatial data is to assign cells to their constituent cellular populations as defined by expression profiles e.g. T-cells, B-cells, malignant cells etc. To date, the dominant paradigm for performing this analysis is to cluster cells based on their expression profile and then perform post-hoc annotation of the clusters based on known markers that delineate cell types^4–9^. As we demonstrate, such a procedure is sub-optimal and new approaches tailored to spatial expression data are required.

The clustering step of most two-step analysis have been performed using methods developed for disaggregated single cell data^4–9^, such as PhenoGraph^10^. A limitation of disaggregate methods is that they ignore spatial information, in particular the identity of neighbouring cells. Neighbourhood information can be highly informative when inferring the cell types, for example if cell types tend to associate due to receptor-ligand signalling. While some recent approaches have begun to address this issue for spatial transcriptomic data^11,12^ using Hidden Markov Random Field (HMRF) models^13,14^, they have thus far only been able to account for an increased affinity of cells of the same type to be neighbours. This autonomous cell type interaction assumption amounts to “smoothing” the assignment of cells in close proximity to originate from the same population. While this is likely a reasonable assumption in many cases, it fails to capture more complex biological scenarios involving non-autonomous signalling between cells of different types. Our first contribution in this work is to develop a generalised HMRF model capable of handling non-autonomous neighbour interactions.

The annotation of clusters to identify their cell type in two-step procedures is typically performed manually. Manual annotation is problematic as it can be subjective and difficult to reproduce^15^. Furthermore, separating the annotation step from clustering means that valuable “prior” information about the expression profiles expected for each cluster are ignored, forcing methods to learn de novo the expression profiles of clusters. While a significant number of methods have been developed to address the cell type annotation problem for disaggregated single cell data^16^, we are not aware of any approaches that incorporate spatial information. Thus, our second contribution in this work is to provide several options for performing joint spatially aware clustering and cell type annotation. As we show in the results, this approach improves clustering accuracy while negating the need to perform laborious and subjective manual cluster annotation.

To address these issues outlined above we have developed a Bayesian model, SpatialSort, to jointly perform spatially aware clustering and cell type annotation. The input into SpatialSort is a cell by marker expression profile matrix, and a graph where edges represent adjacency between pairs of cells in space. To capture spatial dependencies between cells, SpatialSort models cell labels using an HMRF. We account for different propensities of cell types to be neighbours via an interaction matrix with entries indicating the affinity of cell types to neighbour each other. We fit the model using Markov Chain Monte Carlo (MCMC) methods. The output of SpatialSort is a clustering of cells, and (optionally) annotated identities of each cluster. To test the performance of SpatialSort we have conducted benchmarking experiments using synthetic and semi-real datasets. We illustrate the utility of SpatialSort by applying it to real world diffuse large B-cell lymphoma (DLBCL) dataset profiled with MIBI. Our results demonstrate SpatialSort is able to leverage spatial information and prior knowledge of cell type composition to improve clustering and annotation of spatial expression data.

## Results

### Probabilistic spatially aware clustering with SpatialSort

We provide a high level overview of the SpatialSort model and inference procedure here, a more detailed discussion can be found in the Methods sections. A schematic overview of the SpatialSort method is provided in **Fig. 1.** SpatialSort jointly considers cell expression values and neighbourhood spatial structure to perform clustering. To perform unsupervised clustering, SpatialSort requires inputs consisting of a multi-sample marker by cell expression matrix and a list of sample-specific cell location matrices from spatial expression profiling, which is used to identify neighbour cells. Neighbouring cells are defined as cells having a spatial proximity less than a user set threshold in pixels. SpatialSort takes the cell location and neighbour relations to construct sample-specific cell connectivity graphs that link cells that are spatially proximal. To capture the non-random spatial associations between cell types, SpatialSort uses an HMRF to allow cells to influence the cluster assignments of their neighbours (**Supplementary Fig. 1**). As exact Bayesian inference for HMRF models is intractable, SpatialSort uses MCMC sampling to approximate the posterior distribution and estimate model parameters.

**Figure 1:**
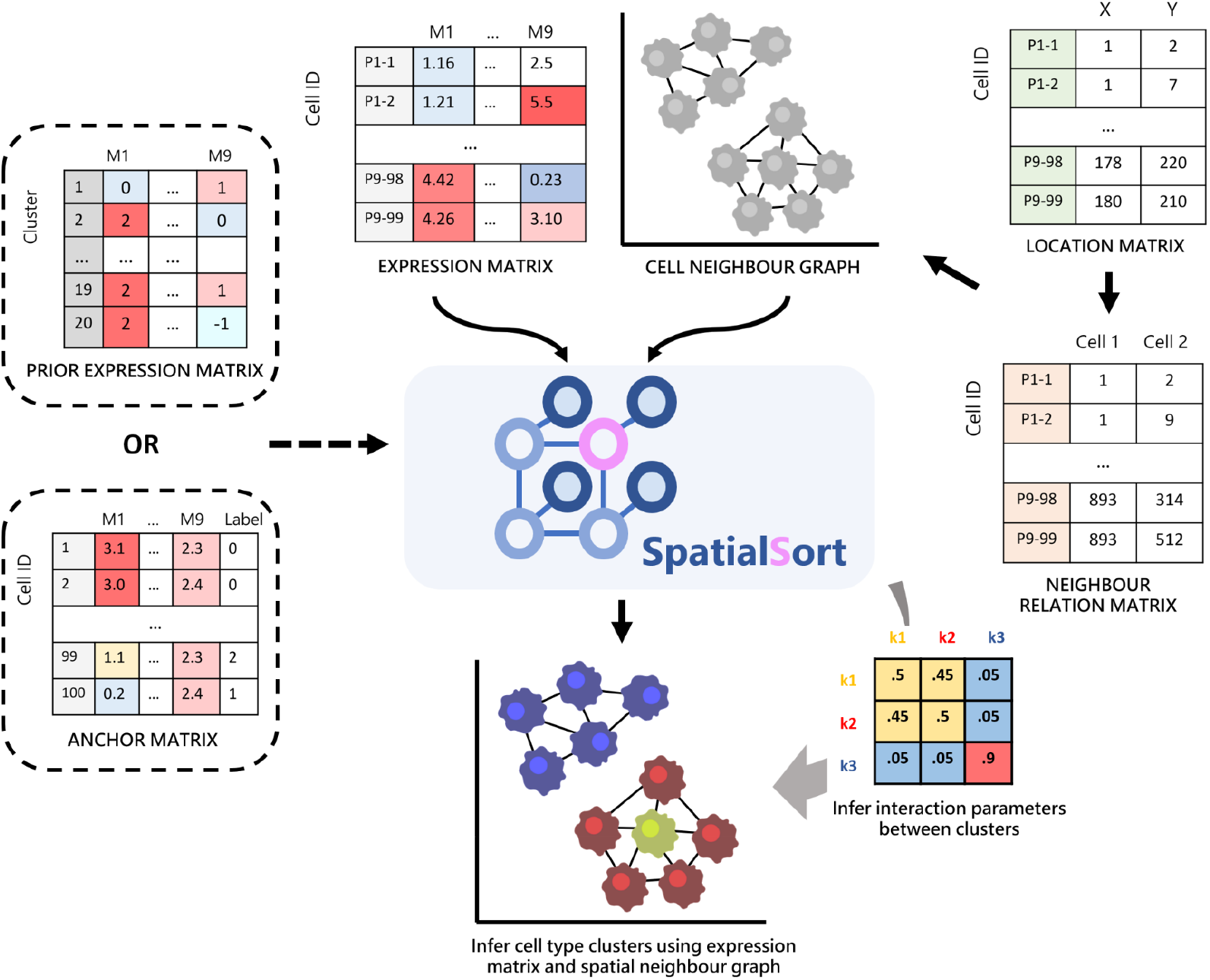
Schematic overview of SpatialSort. SpatialSort requires expression, cell location, and neighbour relation data as inputs. For each patient, a neighbour graph modeled by a MRF is built to represent the spatial context. Using both expression data and spatial structure for inference, SpatialSort jointly infers cluster assignment and the interaction parameter of the HMRF to probabilistically assign each cell to a given cell type cluster in an unsupervised setting. When an expectation of certain cell types or a collection of labeled data is present, a prior expression matrix or an anchor expression matrix can be incorporated to improve clustering or perform label transfer.

SpatialSort can be run in a completely unsupervised way when no prior information is available about cell populations. However, the majority of spatial proteomic studies utilize markers chosen to discriminate among known cell populations. SpatialSort provides two modes for incorporating information about these known populations expression profiles. *Prior* mode takes as an additional input a population by marker matrix, which encodes prior knowledge of the degree of expression per marker in each cell population. *Anchor* mode involves the introduction of anchors cells, which are expression profiles of cells measured by previous assays and assigned to cell populations. Multiple cells from each population may be included in the set of anchors, which can better reflect the variability of expression within the population and aid SpatialSort in inferring the expected variance of marker expressions. We do not model spatial effects for anchor cells, and thus anchor cells can be measured using either disaggregated or spatial technologies. The major constraint for anchors are: **i)** that a reasonable number of overlapping markers are covered in the anchor and the query dataset, **ii)** the anchor dataset is suitably transformed to have expression which match the query dataset. We illustrate the use of anchors derived from disaggregate CyTOF to analyze a MIBI dataset later. Both prior and anchor mode can accommodate the discovery of unknown populations, for prior mode this amounts to specifying clusters with vague priors for all markers and for anchor mode specifying clusters with no anchor cells.

### Modelling non-autonomous cell interactions increases accuracy

We first sought to systematically explore the impact of incorporating spatial information during clustering. To do so we simulated data from the SpatialSort model allowing for non-autonomous cell to cell affinities. To simulate real spatial structure, breadth first search was applied on neighbourhood graphs generated from a previous IMC study^8^ as it maintains the spatial structure of the subset graph. We explored variations of expression values and spatial structure by generating 100 datasets each for two types of HMRF interactions parameters which we refer to as ‘biased’ and ‘uniform’. Biased refers to the condition where cells of the same cluster had a stronger affinity to be grouped together spatially, whereas uniform referred to the case where affinities were sampled from a uniform distribution. We used these datasets to evaluate three variants of the SpatialSort model differing in the number of parameters used to model cell to cell affinities: ‘0p’ - a single fixed parameter for autonomous affinities; ‘1p’ - same as 0p but with the parameter estimated; ‘*K*p’ - one parameters per cluster to reflect autonomous and non-autonomous cell to cell affinities (Methods). We also compared against a Gaussian mixture model (GMM), which is effectively a non-spatial equivalent to SpatialSort.

The results of this analysis are summarised in **Supplementary Figs. 2** and **3** for the biased and uniform datasets respectively. Clustering accuracy was assessed using the V-Measure metric with a value of 1.0 indicating perfect accuracy^17^ (Supplementary Table 1). When comparing methods we applied the Friedman test to see if there were any significant differences in performance between the methods (p-value<0.01) (Supplementary Table 4). If the Friedman test was significant we then applied the post-hoc Nemenyi test with a Bonferroni correction to all pairs of methods to determine which methods showed significantly different performance from each other (p-value < 0.01)^18^(Supplementary Table 7). All statements of significance are with respect to this procedure. All variants of the SpatialSort model significantly outperformed the GMM in our experiments. The simplest SpatialSort model, the 0p model, had a V-measure which was 0.138 higher than the GMM on average for both the biased and unbiased datasets. The *K*p model had significantly better accuracy than both the 0p and 1p models for both biased and uniform datasets. For the biased dataset, the V-measure was on average 0.027 and 0.057 higher for the Kp model when compared to the 0p and 1p models respectively. The performance delta between Kp and simpler spatial models was much larger for the uniform datasets. The Kp model had an average increase of V-measure of 0.112 and 0.093 over the 0p and 1p models respectively.

The increased accuracy of all variants of the SpatialSort model in comparison to the GMM highlights the importance of accounting for spatial structure. The increased delta in performance between the Kp and simpler spatial models supports the notion that explicitly accounting for non-autonomous cell to cell interactions can lead to significant gains in performance when such interactions are present.

### SpatialSort is robust to overlapping expression profiles

We posited that accounting for spatial structure would improve cluster assignment in the case of cells with similar expression profiles. To explore this hypothesis simulated data using the same strategy as the previous synthetic experiment, but varied the degree of overlap in marker expression distributions. Marker expressions were modelled using Gaussian mixtures that were generated using the MixSim R package^19^, which allowed for controllable overlap of simulate expression profiles. We evaluated across 5 different overlaps from 0.025 to 0.125 and varied spatial structure by generating 50 datasets for each overlap under both biased and uniform interaction parameters. For this analysis we consider only the *K*p variant of the SpatialSort model, henceforth referred to as SpatialSort. We compared against GMM as a baseline, and also Phenograph^10^ which is a widely used clustering approach for spatial data. We again applied the Friedman and post-hoc Nemenyi test to assess statistical significance.

Results from this experiment are summarized in **Supplementary Figs. 4** and **5**. SpatialSort significantly outperformed the GMM and Phenograph for all overlap values on both the biased and uniform datasets. The average increase of V-measure for SpatialSort versus GMM ranged from 0.210 to 0.399 and versus Phenograph ranged from 0.128 to 0.492 (Supplementary Tables 2 and 8). The performance of all methods degraded as the degree of overlap in expression profiles increased. However, SpatialSort’s performance was significantly more robust to increasing overlap (Supplementary Tables 5 and 8). This trend held for both biased and uniform datasets. These results support the hypothesis that spatial information can help to more accurately cluster cell types with similar expression profiles.

### Prior information improves accuracy

We next sought to explore the impact of incorporating prior information during clustering. To do so we generated *semi-real* datasets by using real cell expression profiles from a 13-dimensional CyTOF bone marrow mononuclear cell data downloaded from Levine et al^10^. Cell labels for this dataset were obtained by manual gating in a previous study^20^ and used as ground truth for our analysis. Cell neighbourhood graphs and node labels were generated the same way as the synthetic experiments. Expression values were associated with nodes in the graph by assigning a cell from the corresponding cluster in the CyTOF data. We explored variations of clusters and spatial structure by generating 100 datasets for the biased and uniform interaction parameters. The compositions of cell types was similar when simulating data with either of the two types of HMRF interaction parameter settings (**Supplementary Fig. 6a-b**), with the difference in datasets manifesting in the spatial organization of cells (**Fig. 2a-b**). We compared SpatialSort in unsupervised, prior and anchor modes to GMM and Phenograph. We performed principle component analysis (PCA) to reduce the dimensionality of the data to 8 dimensions for GMM, unsupervised SpatialSort and SpatialSort with anchors. No dimensionality reduction was applied when using prior mode for SpatialSort, as specifying prior values of principle components was not a realistic use case. Phenograph was also run without PCA dimensionality reduction, as it applies its own dimensionality reduction.

**Figure 2:**
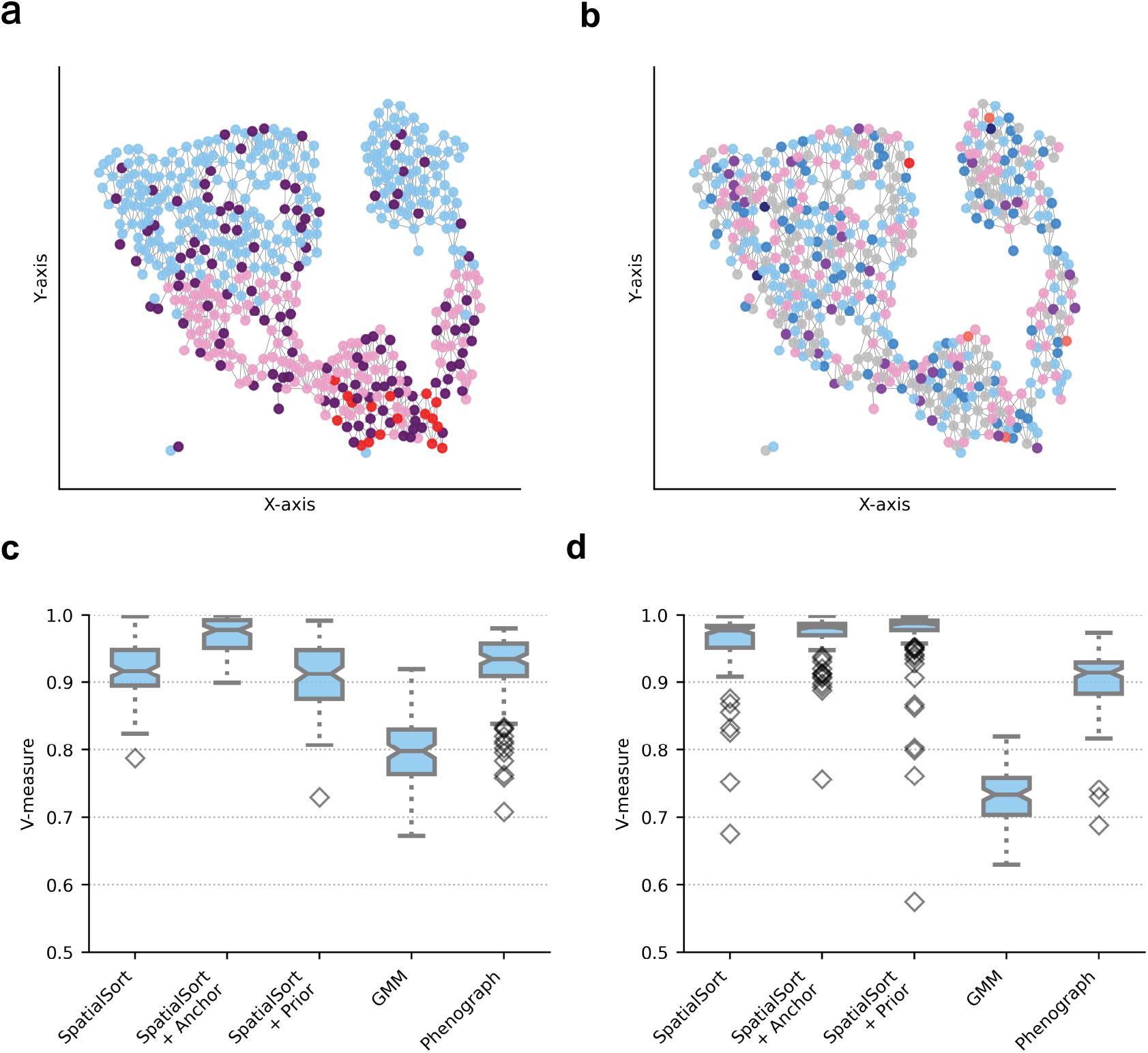
Performance on semi-real spatial CyTOF data. **a)** An example of a spatial neighbour graph of a singular sample in the biased dataset. Nodes indicate a single cell color-coded by cluster assignment. Cells tend to engage in autonomous interactions spatially. **b)** An example of a spatial neighbour graph of a single sample in the uniform dataset as a comparison. Uniform interaction terms render cells to have a random chance of neighbouring any type of cell. **c)** Boxplot of V-measure scores to show clustering accuracy of various methods fitting on 100 semi-real biased datasets and **d)** uniform datasets.

Results of this experiment are summarized in **Fig. 2c-d** and Supplementary Tables 3, 6 and 9. SpatialSort was significantly more accurate than GMM for both biased and uniform datasets using all three modes. The average increase in V-measure for SpatialSort ranged from 0.133 to 0.259. There was no significant difference in performance between SpatialSort in unsupervised mode and Phenograph for the biased dataset, and unsupervised SpatialSort significantly outperformed Phenograph in the uniform dataset. When using prior mode, there was no significant difference between SpatialSort and Phenograph for the biased dataset, and again SpatialSort significantly outperformed Phenograph for the uniform dataset. SpatialSort demonstrated its best performance in the anchor mode. Using anchors SpatialSort outperformed GMM and Phenograph on both the biased dataset and the uniform dataset. SpatialSort in unsupervised mode was significantly outperformed in all cases by both prior and anchor modes. A significant difference in performance was observed between prior and anchor modes in the biased dataset, however it was not observed in the uniform dataset with both methods reporting V-measures near 0.97. These results suggest the including prior or anchors information significantly improves the accuracy of spatially aware clustering.

### Employing anchors to characterize the spatial architecture of DLBCL

To illustrate the real-world utility of SpatialSort we next analysed a MIBI dataset of 116,000 cells from a cohort of 29 patients with DLBCL. For each patient, two regions of interest (ROI) were obtained to address variations in tumour content. We also incorporated the expression data of 128,673 cells from a previously clustered CyTOF assay of the same 29 patients to provide anchors for the characterization of the cellular composition of the tumour micro-environment in the MIBI data. We further subsetted the MIBI and CyTOF data by retaining only marker channels present in both modalities, which were CD45, CD19/PAX5, CD3, CD4, CD8, CD45RO, CD57, CXCR5, PD-1. A linear normalization was applied to scale data from the two modalities to the same expression range, and dimensionality reduction with PCA was applied. We then ran the 0p, 1p and SpatialSort (Kp model) models with anchors to perform label transferring.

The results of this analysis are summarized in **Fig. 3** and **Fig. 4.** With spatial data, we were able to investigate the interaction matrices which indicate the observed frequency of two cell types to be spatially proximal. All three spatial models were able to capture the strong autonomous interaction between B cells (**Fig. 3a-b**, **Supplementary Fig. 7a**) due to the property of DLBCL having substantially higher tumour cell content than cells of other types^21^ (**Supplementary Fig. 8)**. However, we observed a significant difference (p-value=0.00, Pearson chi-squared test) in the cell type distribution estimated by the 0p model compared to the SpatialSort and 1p models (**Fig. 3c-d**, **Supplementary Fig. 7b**).

**Figure 3:**
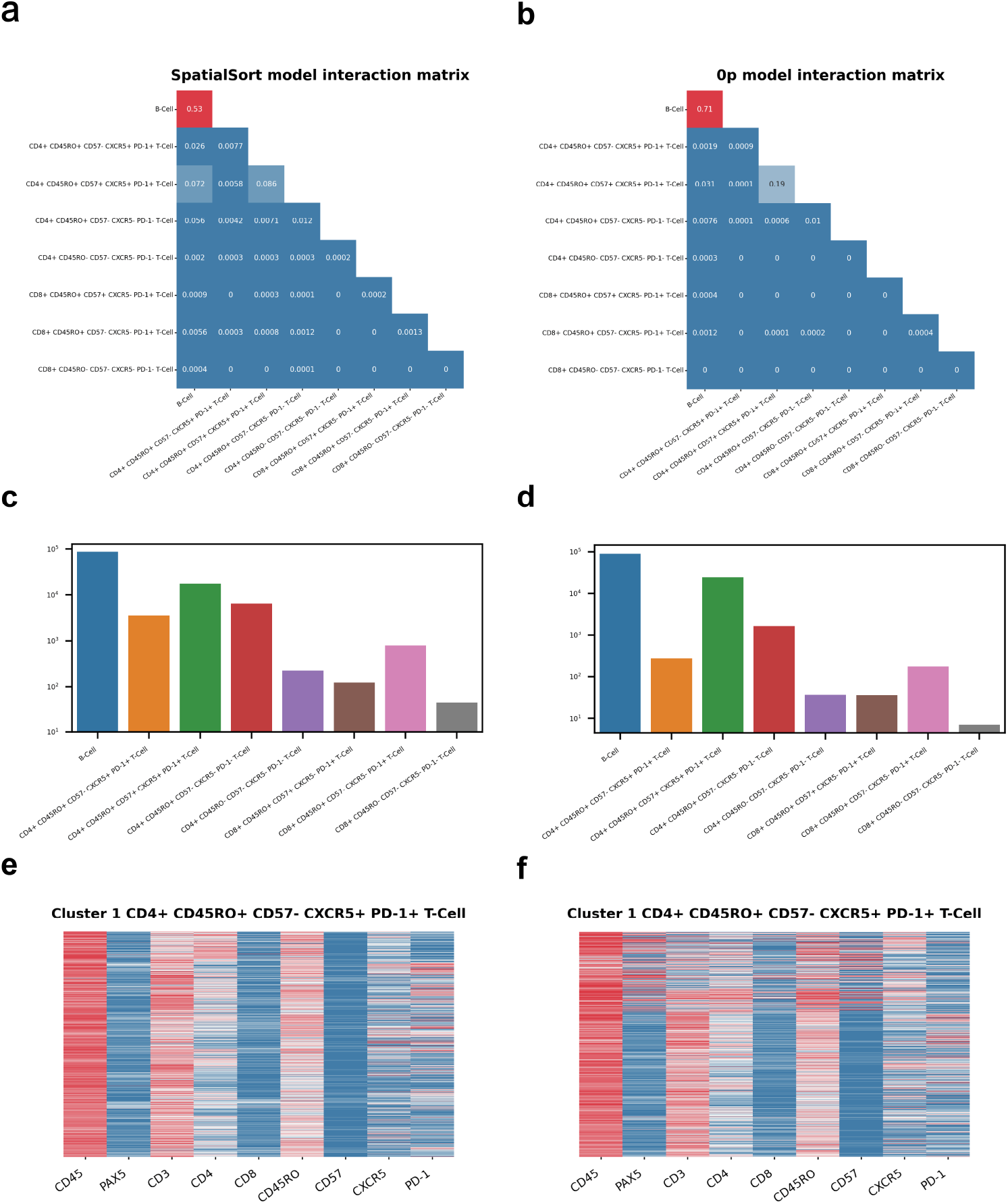
Cell type annotation of DLBCL MIBI data using SpatialSort allows for more effective cell-cell interaction analysis than the 0p model. **a)** The interaction matrix for 29 patients with DLBCL generated using the SpatialSort model. Each cell of the matrix represents the probability distribution of an edge to be between two cell types in the HMRF. An edge represents cells of a cell type to be spatially proximal and interacting with cells of another cell type. **b)** The interaction matrix for 29 patients with DLBCL generated using the 0p model. **c)** The cell type distribution bar graph of the clustering results from using the SpatialSort model. Counts are log-scaled. **d)** The cell type distribution bar graph from using the 0p model. **e)** An exemplar cluster heatmap of a CD4+ CD45RO+ CD57-CXCR5+ PD-1+ T cell from using the SpatialSort model. **f)** A cluster heatmap of the same cell type from the 0p model for comparison.

**Figure 4:**
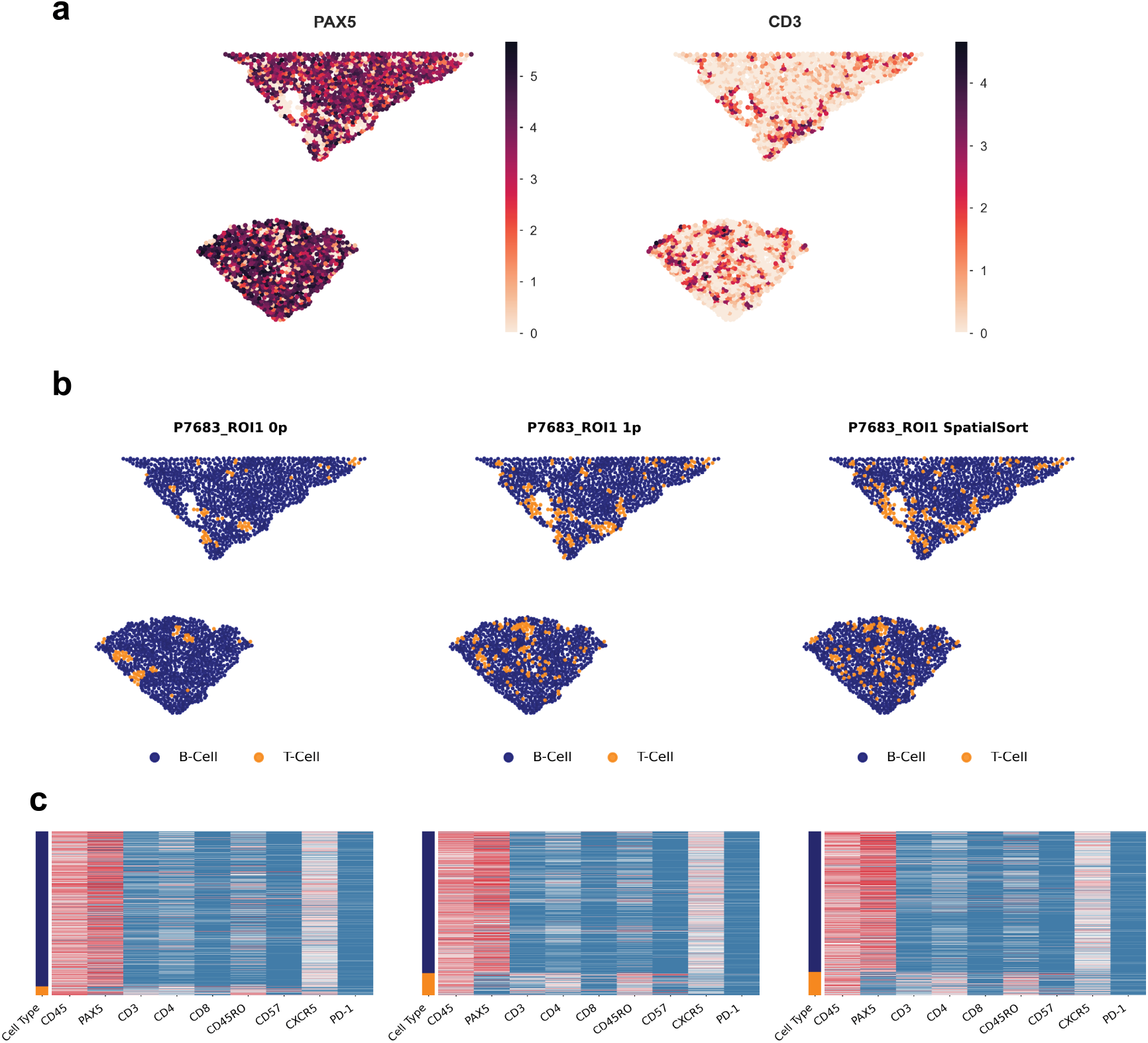
Cellular associations in the spatial organization of DLBCL MIBI data depicted by patient-specific neighbour graphs using SpatialSort. **a)** Spatial distribution of the expression of lymphocyte lineage markers, PAX5 and CD3, across cells in sample P7683. Color represents normalized intensity of expression. **b)** Neighbour graphs of sample P7683 plotted by spatial coordinates. Cells are color-coded by cell type assignment inferred by the 0p model, 1p model, and SpatialSort in anchor mode. **c)** Sample-specific expression heatmaps for sample P7683. Rows are color-coded by cell type in (b).

Visualization with cluster specific heatmaps (**Fig. 3e-f**, **Supplementary Fig. 7c**) revealed some clusters from the 0p model having higher disparity in expression patterns between cells than that of SpatialSort and 1p models. Applying the Davies-Bouldin score^22^, SpatialSort and the 1p model were superior at 1.92 compared to the 0p model at 2.67, with a lower score indicating higher coherence and less noise within clusters. Additionally, visualization of the cellular associations in the spatial structure using patient-specific neighbourhood graphs depicted an over-smoothing effect from the 0p model compared to the 1p and SpatialSort models (**Fig. 4a-c**). An exemplar from sample P7683 illustrates that SpatialSort can more effectively resolve cell types consistent with lineage marker intensities and effectively distinguish between cell types with overlapping expression profiles. Furthermore, these results suggests that the non-random associations between cellular phenotypes in the spatial structure can be more effectively identified when autonomous and non-autonomous interactions are inferred in spatially aware clustering.

## Discussion and conclusions

SpatialSort provides two important advancements over current state of the art methods for analysing spatial protein expression data. First, SpatialSort accounts for potential affinities between non-autonomous cell neighbours while clustering. This more accurately models the underlying biology and improves over the smoothing approach implicit in current HMRF based models^11,12^. Second, SpatialSort provides the ability to incorporate prior information about the expected cellular populations present. This improves upon post-hoc labelling of clusters due to the fact that prior information is directly incorporated while clustering, increasing accuracy.

SpatialSort’s main limitation is computational complexity due to the challenges of posterior inference. The posterior distribution is doubly intractable because not only is the normalization constant of the posterior distribution difficult to evaluate explicitly, as is typical for Bayesian models, but also the likelihood of the HMRF. Previous HMRF based approaches have avoided this issue by using a single autonomous affinity value set manually^11,12^, thus avoiding the need to compute the normalization constant of the HMRF. Our results suggest this limits current HMRF methods to effectively be spatial smoothers. We address this issue using the double Metropolis-Hastings algorithm to approximately sample from the posterior. However, this precludes the possibility of using more computationally efficient approaches such as expectation maximization and variational methods for inference. Despite this, our analysis of real datasets with over 100,000 cells took on average 1.1 minute per sampling iteration or 9 hours to perform an entire run on a personal laptop computer. For extremely large datasets we would suggest downsampling the number of cells based on a breadth first search of the neighbour graphs. Though we have not explored it in this work, there is also significant opportunity for parallelisation across disconnected components of the neighbour graph.

In this work we have primarily focused on the application of SpatialSort to proteomic data. However, there is no reason the model could not be modified to work with transcriptomic data. The key consideration would be that transcriptomic data is typically integer valued in contrast to proteomic data which is continuous. To address this, the user could perform a suitable transformation of the count data to make it continuous as is common in the differential expression literature^23^. An alternative approach we leave for future work would be to replace the Normal emission distribution for the data with discrete distribution such as a Negative-Binomial^24^.

We believe SpatialSort will be a valuable contribution to the spatial expression toolbox for many biologists. It addresses several unmet needs in the field and identifies several novel issues that have thus far been ignored.

## Methods

### A generative model for spatially-aware clustering of expression data

SpatialSort is an instance of a Hidden Markov Random Field (HMRF) model. HMRFs models are defined on an undirected graph *G* = (*E, V*) where *E* is the set of edges in the graph and *V* are the set of vertices or nodes. Because the graph is undirected we assume that E is a set of sets, where elements of E are sets of the form {*u, v*} with *u, v* ∈ *V*.

Let the observed data be denoted by 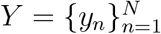, where *N* is the total number of data points and *N* = |*V*|, in the case of SpatialSort a data point is the measured expression profile of a cell. We assume 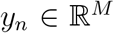 where *M* denotes the number of proteins measured. Each data point *y_n_* has an associate latent variable *x_n_* ∈ {1,…, *K*}, where *K* is the number of clusters or cell populations in the case of SpatialSort. Let 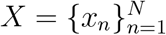 denote the set of all latent cluster allocation variables where each *x_n_* is the label of a node *n* in the graph G. We assume *X* follows a Markov Random Field (MRF) distribution where the value of *x_n_* depends on the values of its immediate neighbours in the graph. We denote the set of neighbours of *x_n_* in *G* by 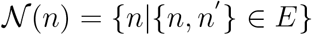. The MRF is governed by *K* × *K* affinity matrix which we denote by *β*. The specification of the priors for the entries of *β* is deferred to the next section where we describe variants of the SpatialSort model. Each cluster *k* has an associated parameter *θ_k_*, which in the case of SpatialSort represents the mean and precision of expression of proteins for cells associated with cluster *k*. Each component of *θ_k_*, denoted *θ_km_*, is assumed to be independent and given a NormalGamma prior distribution. Given *x_n_* and 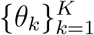 we assume the values of *y_n_* are conditionally independent. The full joint distribution for the model is given in equation 1.

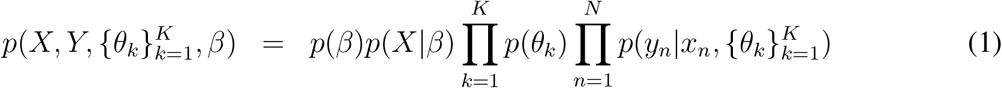

The term *p*(*X*|*β*) describes the MRF component of the joint distribution. The MRF distribution is a product of terms for each edge in the graph. Each term in the product is the exponential of the entry in the matrix *β* corresponding to the identity of contributing edges. The unnormalized form of *p*(*X*|*β*) is given in equation 2.

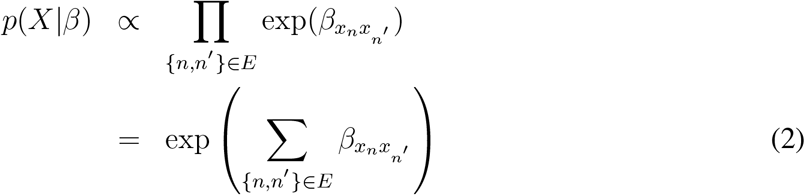

The normalization constant *Z*(*β*) of *p*(*X*|*β*) can be found by summing over all possible values of 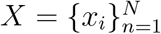, which is intractable for all but small values of *N*. As we discuss later this poses an inferential challenge when updating *β*.

Thus the full hierarchical model, except for the specification of *β*, is as follows.

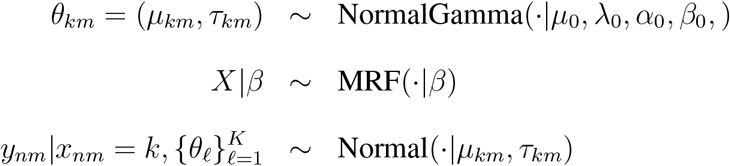

The model can be trivially extended to multiple samples or regions of interest by treating each new sample as separate connected components of the MRF graph.

### Specifying the affinity matrix

The affinity matrix *β* is assumed to be symmetric, thus there are up to 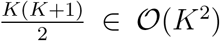 free parameters that need to be specified. In practice, it neither computationally feasible nor statistically efficient to treat all entries of *β* as free parameters. Here we discuss several parameterizations of *β* which lead to different variants of the SpatialSort model.

The simplest and most commonly employed parameterizations of *β* is to use a single value, *β^s^*, which is shared across all diagonal entries and setting the off diagonals to 0 i.e. *β_kk_* = *β^s^* and *β_kl_* = 0 for *k* ≠ *l*. This simple model, often referred to as as the Potts model, captures affinities of cells of the same type and assumes that they all have the same strength. Due to the intractability of the normalization constant *Z*(*β*) of *p*(*X*|*β*), it is common to fix *β^s^*. We refer to the variants of SpatialSort with *β^s^* fixed as the 0p and with *β^s^* estimated as the 1p model. For the 1p model we assign *β^s^* a Uniform(0,1) prior. For the 0p model we fix β^s^ to 0.5 for all analyses performed in this work.

The limitation of the standard Potts model, is the inability to capture affinities between clusters (cell populations) of different types. To address this we consider a richer parameterization of *β* which allows for variable strengths of autonomous interactions, and allows for non-autonomous interactions. We refer to this model as the *K*p model, as there are *K* parameters which need to be estimated. In the *K*p model the diagonals of *β* are set to 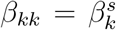 which accounts for variable affinities for autonomous interactions. We define 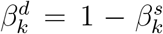 and let 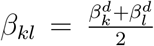 for the off diagonal terms to capture non-autonomous interactions. We assign a Uniform(0,1) prior to 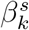.

### Incorporating prior knowledge into clustering

The incorporation of prior knowledge of the marker proteins can be applied to improve clustering accuracy. A quaternary coded *K* by *M* prior expression matrix can serve as an input parameter of SpatialSort, where each row is a prior belief of the marker intensities for a cluster. Through coding values from 0 to 2, the mean parameter *μ_km_* of *θ_km_* is then translated to the 25th, 50th, and 75th percentiles for each marker expression of *Y*. The value −1 is a special case which translates to a zero mean coupled with an high variance, which occurs in the case when we do not have prior knowledge on the expression of markers.

Another approach is to leverage previously annotated cell types and anchor clusters to specific expression profiles. The introduced cells are referred to as anchors, as they are observed variables influencing the updates of cell cluster assignments and strongly anchor clusters to a specific expression signature profile. Anchors have a fixed cluster assignment and do not contribute to the HMRF graph. The anchors act to specify the distribution parameters of their associated clusters. This approach improves the accuracy of clustering and allows for label transfer between disaggregate and spatial datasets.

### Inference of latent cluster labels and cell-cell interactions

Inference on *X* and *β* constitutes of computing the (marginalized) posterior distribution, which can be formulated as:

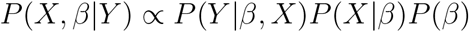

Closed-form solutions are intractable due to the complexity of the model, instead we employ Markov Chain Monte Carlo sampling methods to approximate the posterior distribution. Cell labels *x_n_* are sampled through a collapsed Gibbs sampler (CGS). The interaction parameters *β* are sampled via a Double Metropolis-Hastings (DMH) sampler^25^. Detailed information about the CGS and DMH steps are described in the **Supplementary Note**. One full iteration of the inference algorithm perform five updates of *β* using the DMH algorithm and one update of *X* using Gibbs sampling.

### Obtaining point estimates of the MCMC trace

Given the approximated posterior distribution of *X* and *β* through sampling, referred to as the MCMC trace, we summarize the posterior by deriving point estimates for downstream analysis.

To derive a point estimate for *X*, we construct a distance matrix using Hamming distance and apply hierarchical clustering. For all experiments on synthetic and real datasets, we ran SpatialSort for 500 iterations. A burn-in portion of half the MCMC trace is removed as standard practice. For unsupervised clustering, we optimize the Maximization of Posterior Expected Adjusted Rand (MPEAR)^26^ criterion which yields a sequence of consensus class labels given the MCMC trace. In anchor mode, we do not optimize MPEAR, instead we use the last sample given the trace has reached convergence.

### Preprocessing

For the semi-real dataset experiments, a 13-dimensional CyTOF dataset of bone marrow mononuclear cells were downloaded from Levine *et al*. Cells without labels from gating were discarded. An arcsin transformation was applied to normalize the dataset. Dimensional reduction with PCA was performed on the markers for unsupervised clustering, anchor mode, and GMM.

For the real-world DLBCL dataset experiments, CyTOF DLBCL datasets were normalized by marker against a spike-in control to account for machine drift and batch effects in staining. This dataset was then normalized by a hyperbolic arcsin function. MIBI DLBCL datasets were also normalized by a hyperbolic arcsin function and divided by 10 to reduce expression intensity to the same scale as CyTOF. As there were no common B cell lineage marker between CyTOF and MIBI, CD19 and PAX5 were treated as equivalent. In the anchor experiments, spatially aware downsampling through breadth first search was performed on the MIBI data to 2000 cells per sample. Addition subsetting was done on both CyTOF and MIBI datasets to retain only overlapping markers: CD45, CD19/PAX5, CD3, CD4, CD8, CD45RO, CD57, CXCR5, PD-1. Dimensional reduction with PCA was performed on the common cell type lineage markers between the two modalities. The top six principal components were used as input for label transferring.

### Benchmarking

For all forward simulations, Gaussian mixture simulations and semi-real simulations, we applied GMM as a benchmarking method using the GaussianMixture function from the scikit-learn package version 0.24.2^27^. The number of components for GMM were set to the same number of clusters as were set for SpatialSort. For the latter two simulations, we additionally applied Phenograph version 1.5.7^10^ with default parameters for benchmarking. Clustering accuracy was assessed using the V-Measure metric which is a harmonic mean between completeness and homogeneity^17^.

## Declaration

### Ethics approval and consent to participate

All samples were obtained with informed consent and according to protocols approved by the BCCA Research Ethics Board.

### Consent for publication

All patients provided written consent for publication.

### Availability of data and materials

The SpatialSort Python package is available on Github at: https://github.com/Roth-Lab/SpatialSort under the MIT license. Raw data for all the experiments used in this article have been deposited in Zenodo with DOI: https://doi.org/10.5281/zenodo.6909419.

### Authors’ Contributions

AR and AW conceived the study design. EL, KC, ABC, and AR designed the statistical method. EL, MN, XW, CH, AG carried out the experiments. EL, KC implemented the software. EL, MN, XW performed the data processing, analysis, and simulations. All authors read and approved the final manuscript.

## Acknowledgements

We provide a detailed listing of the members of the IMAXT Consortium: Mohammad Al Sa’d, Hamid Raza Ali, Martina Alini, Samuel Aparicio, Heather Ashmore, Thomas Ashmore, Shankar Balasubramanian, Giorgia Battistoni, Robby Becker, Bernd Bodenmiller, Edward S Boyden, Dario Bressan, Alejandra Bruna, Marcel Burger, Carlos Caldas, Maurizio Callari, Ian Gordon Cannell, Nick Chornay, Ali Dariush, Lauren Deighton, Lauren Deighton, Khanh N Dinh, Yaniv Eyal-Lubling, Jean Fan, Atefeh Fatemi, Debarati Ghosh, Eduardo A GonzÀ¡lez-Solares, Wendy Greenwood, Flaminia Grimaldi, Gregory J Hannon, Owen Harris, Suvi Harris, Cristina Jauset, Johanna A Joyce, Tatjana Kovačević, Laura Kuett, Russell Kunes, Daniel Lai, Emma Laks, Hsuan Lee, Giulia Lerda, Yangguang Li, Jack Lovell, Yangning Lu, John Marioni, Andrew McPherson, Neil Millar, Alireza Molaeinezhad, Claire M Mulvey, João CF Nogueira, Fiona Nugent, Ciara H O’Flanagan, Marta Paez Ribes, Isabella Pearsall, Sarah Pearsall, Brett Pryor, Fatime Qosaj, Clare Rebbeck, Andrew Roth, Oscar M Rueda, Teresa Ruiz, Kirsty Sawicka, Leonardo A Sepúlveda, Sohrab P Shah, Abigail Shea, Anubhav Sinha, Austin Smith, Simon Tavaré, Ignacio Vázquez-García, Sara Lisa Vogl, Nicholas A Walton, Spencer S Watson, Joanna Weselak, Tristan Whitmarsh, Jonas Windhager, Ruihan Zhang, Chi Zhang, Pu Zheng & Xiaowei Zhuang.

