## Supplementary Methods and Figures for "SpatialSort: A Bayesian Model for Clustering and Cell Population Annotation of Spatial Proteomics Data"

**Bibliography** 10

**2 Supplementary Figures** **11**

**3 Supplementary Tables** **18**

### 1 Supplementary Note

#### 1.1 Mean and Precision: Normal-Gamma Distribution

We assume the cluster parameters $\theta_{km}$ are conditionally independent for both the $k$ and $m$ indices. The parameter $\theta_{km}$ can be written as $(\mu_{km}, \tau_{km})$ on which we assume a Normal-Gamma distribution. For ease of notation we drop the subscripts $k$ and $m$ in this section, with the understanding the same procedure is used for prior specification all $\theta_{km}$.

The Normal-Gamma distribution is conjugate to the Gaussian distribution with unknown mean $\mu$ and precision $\tau$. The Normal-Gamma distribution is defined as follows:

$$\begin{aligned} \tau &\sim \text{Gamma}(\alpha_0, \beta_0) \\ \mu | \tau, \lambda_0 &\sim \text{Gaussian}(\mu_0, \sqrt{1/(\lambda_0 \cdot \tau)}) \end{aligned}$$

in which $\alpha_0$ is the shape parameter and $\beta_0$ is the rate parameter for the Gamma distribution.

We set $\lambda_0 = 0.1$. The parameters for the $\alpha_0$ and $\beta_0$ are are selected using the variance of the expression data and parameter searching as described below.

$$\begin{aligned} \mu_\tau &= 1/(\lambda_0 \cdot s_Y^2) \\ \sigma_\tau^2 &= \begin{cases} 1 & \text{if uncertainty is low} \\ 100 & \text{if uncertainty is high} \end{cases} \\ \alpha_0 &= \mu_\tau \cdot \beta_0 \\ \beta_0 &= \mu_\tau / \sigma_\tau^2 \end{aligned}$$

where $s_Y^2$ is the sample variance of $Y$.

The parameter $\mu_0$ is determined by the prior matrix given as a user input. The input prior matrix has dimensions $K$ clusters by $M$ marker and is discrete with quaternary codes. For each value of $k$ and $m$, $\mu_0$ is translated to the 25th, 50th, and 75th percentiles for each marker expression if the prior matrix has codes 0, 1, 2 respectively. $-1$ is a special case which means a value of 0. Having a code of 2 for a particular marker of a cluster indicates prior knowledge of a high expression value. The values 1 and 0 follow the same idea and represents a user's belief of middle and low to no expression respectively. The code "$-1$" is only used when coupled with a extremely uncertain precision which occurs in the case when we do not have prior knowledge on the expression of markers.

#### 1.2 Updating X: Collapsed Gibbs Sampling

In the updates of the labels $X$ of cells, we exploit the properties of conjugacy. Recall that the observed expression data $y_{nm}$ given $x_n = k$ is distributed according to a Gaussian distribution with parameters $(\mu_{km}, \sigma^2_{km})$. We model the parameters to have a Normal-Gamma distribution which is the conjugate prior of the Gaussian distribution. Let us formulate the computation of the conditional distribution used by our Gibbs algorithm as,

$$\begin{aligned} P(x_n, \{\theta_k\}_{k=1}^K | Y, X_{-n}, \beta) &\propto P(Y|\{\theta_k\}_{k=1}^K, X)P(X|\beta)P(\theta) \\ &= \Big(\Pi_{k\in K}\Big(\Pi_{j\in I_k} P(Y_j|\theta_k)\Big)P(\theta_k)\Big)P(X|\beta) \end{aligned}$$

where $I_k = \{n : x_n = k\}$ and $X_{-n} = \{x_j : j \neq n\}$.

We calculate likelihood as a product of probabilities per cluster $k$. We can further expand this equation to incorporate conjugacy to form a collapsed likelihood. The derivation is as follows:

$$\begin{aligned} P(x_n|Y, X_{-n}, \beta) &\propto \int \Pi_{k\in K}\Big(\Pi_{j\in I_k} P(Y_j|\theta_k)\Big)P(\theta_k)P(X|\beta)d\theta_k \\ &= \Pi_{k\in K}\left\{\int \Pi_{j\in I_k} P(Y_j|\theta_k)P(\theta_k)d\theta_k\right\}P(x_n|X_{-n}, \beta) \\ &= \Pi_{k\in K} Z_k(\mu_0, \lambda_0, \alpha_0, \beta_0, Y_k)P(x_n|X_{-n}, \beta) \end{aligned}$$

where $Y_k = \{y_n : n \in I_k\}$. Thus we sample from $P(x_n|Y, X_{-n}, \beta)$ to perform Gibbs updates of $x_n$, where $Z_k(\mu_0, \lambda_0, \alpha_0, \beta_0, Y_k)$ is specified in the following section. To update all values of $X$ we visit nodes in random order at each iteration, sampling new values from $P(x_n|Y, X_{-n}, \beta)$.

Using conjugacy, we can derive a closed form solution for $Z_k(\mu_0, \lambda_0, \alpha_0, \beta_0, Y_k)$, which is the marginal likelihood of the Normal-Gamma distribution.

$$Z_k(\mu_0, \lambda_0, \alpha_0, \beta_0, Y_k) = \frac{\Gamma(\bar{\alpha}_k)}{\bar{\beta}_k^{\bar{\alpha}_k}}\left(\frac{2\pi}{\bar{\lambda}_k}\right)^{1/2}$$

where

$$
\begin{aligned}
n_k &= |I_k| \\
\bar{\lambda}_k &= \lambda_0 + n_k \\
\bar{\alpha}_k &= \alpha_0 + n_k/2 \\
\bar{\beta}_k &= \beta_0 + \frac{1}{2}\sum_{i \in I_k}(y_i - \bar{y}_k)^2 + \frac{\lambda_0 n_k (\bar{y}_k - \mu_0)^2}{2(\lambda_0 + n_k)}
\end{aligned}
$$

in which $n_k$ refers to the number of cells in cluster $k$ and $\bar{y}_k = \frac{1}{n_k}\sum_{i \in I_k} y_i$ is the mean of expression for cells assigned to cluster $k$.

#### 1.3 Updating Beta: Double Metropolis Hastings

To estimate the cell-cell interactions term, $\beta$, we use Double Metropolis Hastings (DMH) algorithm[1]. DMH is an approximate sampling scheme which can be applied when the likelihood is intractable, a so called doubly intractable inference problem. For our model computing the normalization constant or partition function of the HMRF term would be required to implement a regular MH update. Computing the partition function is intractable as it requires enumerating all possible ways to label the MRF graph. DMH performs an inner round of MH sampling to cancel the partition functions out in the accept-reject ratio.

Our implementation of DMH is as follows. Given the current interaction term $\beta$, the algorithm proceeds in steps:

1. Sample $\beta'$ from the prior $P(\beta)$
2. Generate $u \sim P^{(m)}(\cdot|x, \beta')$ where $u$ denotes an auxiliary variable, $P^{(m)}(y|x)$ is the transition probability from $x$ to $y$, and $m$ denotes the number of Metropolis Hastings iterations used to generate $u$
3. Accept $\beta'$ with probability
$$\frac{f(u|\beta_0)f(x|\beta')}{f(x|\beta_0)f(u|\beta')}$$

We perform $m = 5$ iterations of the inner MH algorithm in step 2 for each iteration of the DMH algorithm.

#### 1.4 Forward Simulating Synthetic Spatial Data

Generating synthetic datasets that are forward simulated from our proposed model serves to evaluate the ability of the inference engine to recover the known true parameters. The simulation of the data starts with the sampling of the interaction matrix for the SpatialSort model, the mean and variance of the expression matrix. The hyperparameters chosen for this specific experiment are detailed in Algorithm 1. For each sample, we simulate a neighbourhood graph by subsetting a part of a real breast cancer imaging mass cytometry topology[2] using breadth first search starting at a random location. The labels of the cells in the topology are then forward simulated from the MRF model using the sampled beta matrix. Expression values for each label are sampled from a Gaussian distribution with parameters from the previously sampled means and variances.

Sample $\beta_k^s \sim Uniform(0, 1)$ for $K$ clusters
Sample $\mu \sim Gaussian(0, 1)$ for $M$ markers of $K$ clusters
Sample $\sigma^2 \sim Gamma(1, 1)$ for $M$ markers of $K$ clusters
**for** *p=1 to P* **do**
    Simulate a topology by Breadth First Search through a real topology structure for sample $p$
    Sample $x_p \sim MRF(\beta)$
    Sample $y_p \sim Gaussian(\mu_{x_p}, \sigma^2_{x_p})$
**end**

**Algorithm 1:** Forward Data Simulation

Using Algorithm 1, we simulated expression data matrices with dimensions 500 cells by 20 protein markers. Each matrix includes a dataset of 10 samples. The number of Gibbs sampling iterations performed for sampling cell labels was set at 5000.

A variant of the algorithm was also used to generate another synthetic dataset with the same parameters described in the previous paragraph. The only change made was to the first line of Algorithm 1 where an extra step was added. The extra step is to swap the autonomous interaction affinity and the non-autonomous interaction affinity if the latter is greater than the former. This step ensures that cells that are of the same type prefer to be with each other than of another type.

For all synthetic and semi-real experiments performed in this chapter, we generate datasets from both two conditions of interest: having random beta affinity values sampled from Uniform(0,1) or having beta affinity values with a stronger affinity for same-same interactions.

#### 1.5 Simulating Gaussian Mixtures Spatial Data

For the MixSim experiments we simulated data as in the previous section, with the exception of the cluster parameters. Cluster parameters were simulated using the MixSim R package[3] which provides a method of simulating Gaussian mixtures with a defined level of overlap. We simulated expression data matrices with dimensions 250 cells by 20 protein markers.

#### 1.6 Simulating Semi-Real Spatial Data

Semi-real datasets are composed of two main elements: (1) expression data from non-spatial expression profiling and (2) simulated topology with sampled labels. The source of expression data used here is by Levine, et al. It is a 13 dimensional CyTOF dataset of a single patient. The 13 surface markers are: CD45, CD45RA, CD19, CD11b, CD4, CD8, CD34, CD20, CD33, CD123, CD38, CD90, and CD3. We used a subset (49%) of the dataset that consists of 81,747 cells of 24 assigned cell type labels from manual gating. The other half of the dataset was not labeled and was not used. The data available was arcsin transformed and no further modification was done to the expression data prior to inference.

The simulation of the data is detailed in Algorithm 2.

Sample $\beta_k^s \sim Uniform(0,1)$ for $K$ clusters
**for** *p=1 to P* **do**
  Simulate a topology by Breadth First Search through a real topology structure for sample $p$
  Sample $x_p \sim MRF(\beta)$
**end**
**for** *k=1 to K* **do**
  Assign mass cytometry expression of the cells labeled $k$ in the non-spatial data to the cells labeled $k$ in the simulated topology
**end**

**Algorithm 2:** Semi-Real Data Simulation

Through the use of Algorithm 2, we generated expression data matrices with dimensions 500 cells by 13 protein markers for 10 samples. For inference, we constructed a prior expression matrix by searching for the markers in the public human protein databases.

#### 1.7 Tissue Staining

4-µm-thick tissue sections from FFPE tissue blocks were cut onto MIBI slides (IONpath catalog number #567001. MIBI is performed by staining tissue with a panel of metal-labeled antibodies and then imaging the tissue using time-of-flight secondary ion mass spectrometry (ToF-SIMS). The slides are stained using IONpath's MIBI protocol. In summary, slides were baked at 70°C for 20 minutes, loaded onto a Histo-Tek SL Slide Stainer (Sakura), deparaffinized with xylene and then rehydrated through successive washes of reagent alcohol (100% decreasing to 70%, Sigma Aldrich) and MIBI diH2O (IONpath). The slides were transferred to a PT Module (Thermo Scientific) and heated to 97°C for 40 minutes in an epitope retrieval buffer (Target Retrieval Solution, pH 9, DAKO Agilent). The slides were blocked with 5% donkey serum (Sigma Aldrich) in TBS-Tween (IONpath). The antibody panel was made by diluting antibody conjugates in the blocking buffer and filtering using a centrifugal filter, 0.1 µm PVDF membrane (Ultrafree-MC, Merck Millipore). The slides were stained overnight at 4°C in a humidity chamber. Slides were loaded into the slide stainer and washed three times with TBS-Tween, fixed for five minutes in 2% glutaraldehyde (Electron Microscopy Sciences), washed three times in 100 mM Tris pH 8.5 (IONpath), twice in MIBI diH2O, and then dehydrated through a series of reagent alcohol washes (70% increasing to 100%). The stained slides were stored in a vacuum cabinet for at least 1 hour prior to loading into the MIBIscope.

#### 1.8 Multiplexed Ion Beam Imaging

The MIBIscope is a dynamic secondary ion mass spectrometer (SIMS) with a time of flight (ToF) mass analyzer. Field of view selection was performed via a point and click interface using the provided H&E images for guidance. The FOVs were confirmed using a secondary electron detector (SED) that shows a live image of the topography of the tissue. FOVs were 800 µm by 800 µm in size and imaged at a resolution of 2048 x 2048 pixels (0.39 µm per pixel). Once the coordinates of all the FOVs for a slide were selected, the MIBI run was executed without any additional user input. The MIBIscope rasters a primary ion beam across the tissue, liberating secondary ions from the metal-conjugated antibodies introduced by staining the tissue. An electrostatic analyzer (ESA) within the MIBIscope acts as an energy filter, significantly biasing the ions detected towards monatomic species and reducing the transmission of polyatomics (hydrides, oxides, organics). The masses of the secondary ions are determined using the orthogonal time of flight mass spectrometer and the detected species were assigned to target biomolecules given the known isotopic label of each antibody. A 33-plex MIBItiff data file is created for each FOV by integrating the mass peaks and correcting for background and mass interferences.

#### 1.9 Cell Segmentation

The MIBItiff data is reviewable using MIBItracker, an interactive, cloud-based data visualization tool. Cell segmentation takes advantage of the multiplexed nature of MIBI data by combining the nuclear dsDNA signal with cytoplasmic and membrane markers to accurately define the seed points and boundaries in tissue images. Initial cell predictions are made using a deep learning-based object-detection model that has been trained on previously segmented MIBIscope data. Boundaries were manually reviewed by visualizing nuclear, membrane, and cytoplasmic signals. As needed, cell segmentations were refined by adding, removing, or moving watershed seed points to ensure cell instances in a given image were accounted for. The result is an image in which each pixel is either assigned an integer value corresponding to a unique cell instance or 0, indicating a cell is not present.

##### 1.10 Cell-of-Origin Algorithm

Cell-of-Origin (COO) for the DLBCL dataset was assigned by Nanostring assay as per Scott *et al*[4]. This particular paper refers to assay as performed on FFPET tissues, but is applied also to RNA from fresh tissues. The particular codeset has been updated from the original Lymph2Cx assay to the DLBCL90 assay[5].

##### 1.11 Computational Performance

The complexity of SpatialSort largely depends on the the DMH step in the updates of the beta interaction term. Time complexity of SpatialSort in terms of big O is $O(K^2PNEtT)$, where $K$ is the number of clusters, $P$ is the number of samples, $N$ is the number of cells, $E$ is the number of edges, $t$ is the number of iterations for DMH, and $T$ is the number of total iterations. Using the Python package Lineprofiler, we profiled the run time of each major parameter updating function. The ratio between updating X and updating beta is 0.43/0.57. The total run time for fitting the model to the forward simulations, spatial Gaussian mixture datasets, and semi-real datasets took on average 0.04 minutes per iteration for between 2500 cells to 5000 cells, and the real dataset took around 1.1 minute per sampling iteration for 116,000 cells on a personal laptop computer.

### 2 Supplementary Figures

**a**

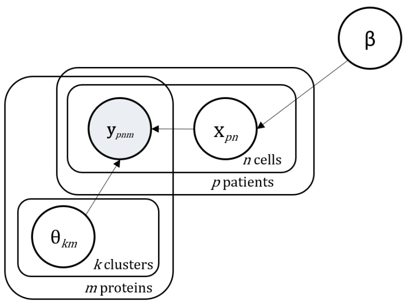

**b**

| Variable | Represents | Distribution |
|---|---|---|
| $y_{pnm}$ | Marker m expression of cell n of patient p | Gaussian |
| $x_{pn}$ | Cell type label of cell n of patient p | Modified Potts |
| $\theta_{km}$ | Combined mean and variance of marker m expression of cluster k | Normal-Gamma |
| β | Cell-cell interaction parameter | Beta |

**Supplementary Figure 1**: **a)** Probabilistic graphical model and **b)** prior distributions of SpatialSort. $y_{pnm}$ represents the expression of marker $m$ in cell $n$ of sample $p$, following a Gaussian distribution. $\theta_{km}$ denotes the combination of mean and precision parameters of marker $m$ in cluster $k$, distributed according to a Normal-Gamma. $x_{pn}$ denotes the label of cell $n$ in sample $p$, following a Modified-Potts distribution with a latent interaction term, $\beta$, which is beta distributed.

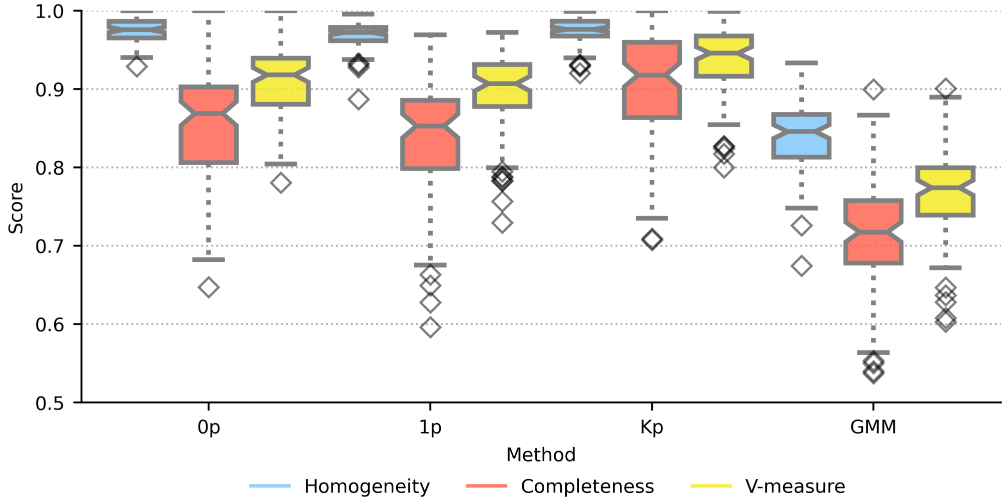

**Supplementary Figure 2**: Comparison of performance on model fitting on forward simulated datasets for biased datasets. Different methods applied to fit the datasets are shown on the x-axis. 0p indicates the Potts model, 1p and $K$p are different parameterizations of the Potts model, GMM indicates the Gaussian mixture model. Scores of performance metrics are shown on the y-axis. Performance was evaluated using three metrics: homogeneity, completeness, and V-measure, color coded according to the legend.

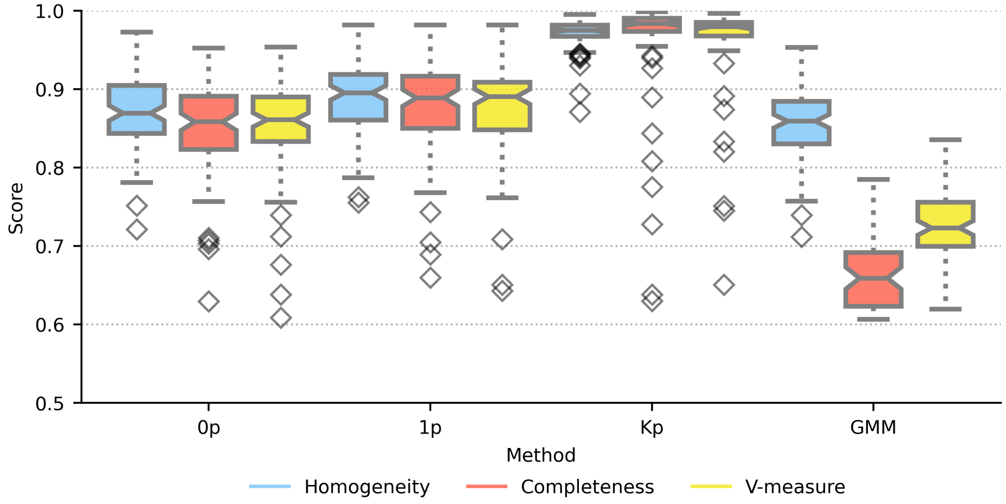

**Supplementary Figure 3**: Comparison of performance on model fitting on forward simulated datasets for uniform datasets. Different methods applied to fit the datasets are shown on the x-axis. 0p indicates the Potts model, 1p and $K$p are different parameterizations of the Potts model, GMM indicates the Gaussian mixture model. Scores of performance metrics are shown on the y-axis. Performance was evaluated using three metrics: homogeneity, completeness, and V-measure, color coded according to the legend.

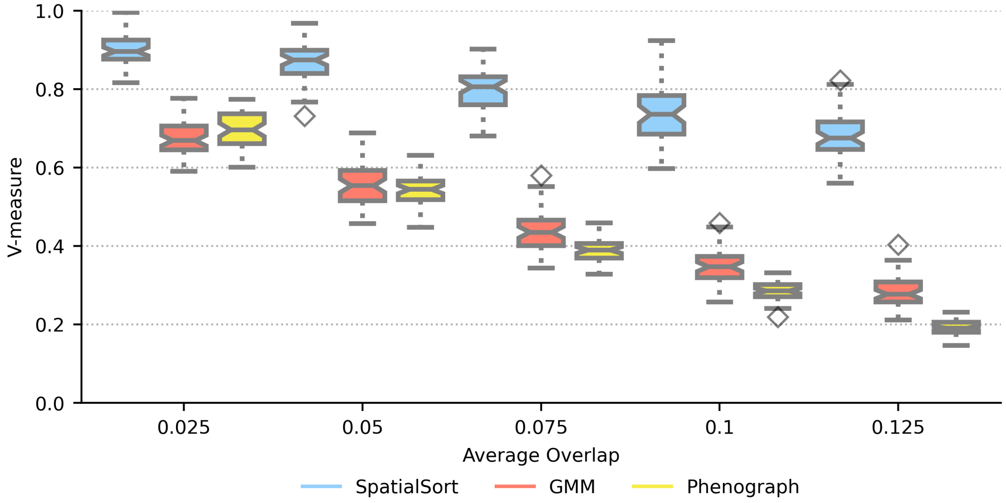

**Supplementary Figure 4**: Comparison of performance on model fitting on spatial Gaussian mixture datasets for biased datasets. The average overlap of spatial Gaussian mixture datasets are shown on the x-axis. Scores of performance using the V-measure metric are shown on the y-axis. Each dataset with different average overlap was fit by three different methods: SpatialSort, GMM, and Phenograph.

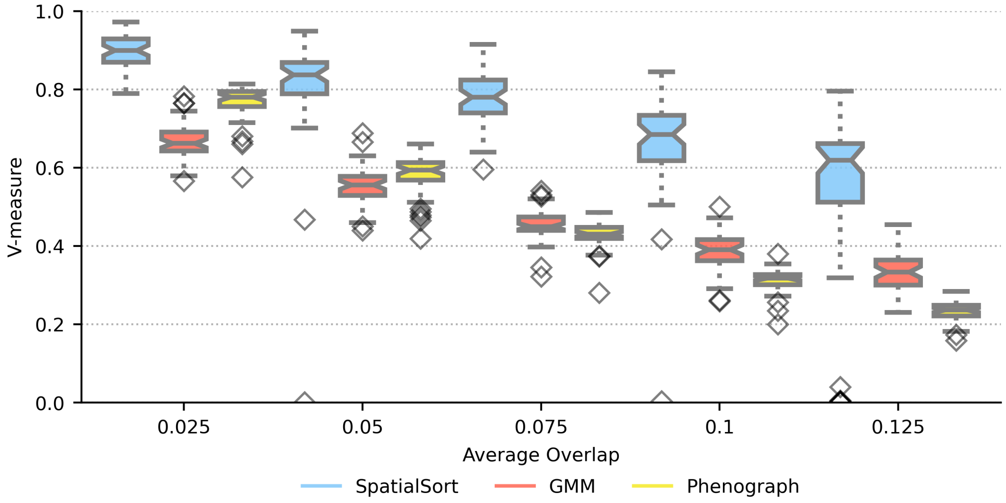

**Supplementary Figure 5**: Comparison of performance on model fitting on spatial Gaussian mixture datasets for uniform datasets. The average overlap of spatial Gaussian mixture datasets are shown on the x-axis. Scores of performance using the V-measure metric are shown on the y-axis. Each dataset with different average overlap was fit by three different methods: SpatialSort, GMM, and Phenograph.

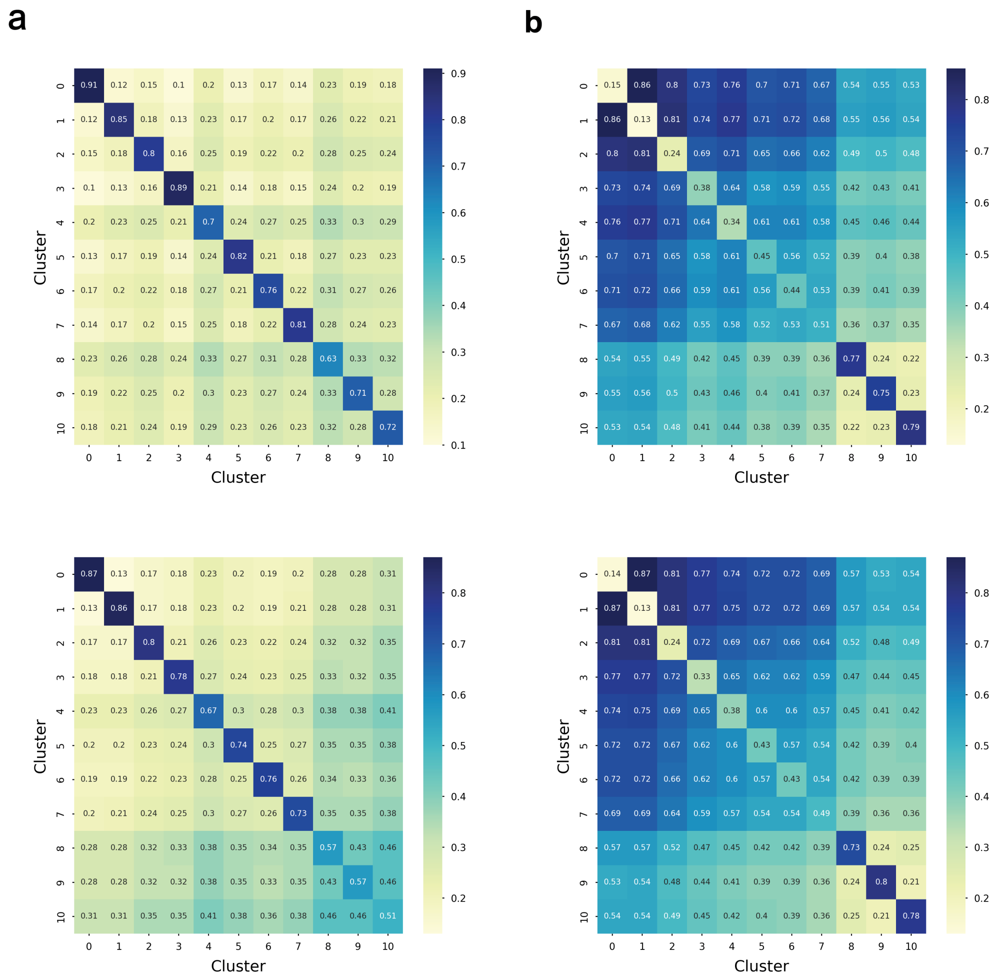

**Supplementary Figure 6**: **a)** Above is the inferred $\beta$ interaction matrix for a semi-real biased dataset, below is the ground truth $\beta$ interaction matrix. **b)** Above is the inferred $\beta$ interaction matrix for a semi-real uniform dataset, below is the ground truth $\beta$ interaction matrix. Axes represent the cluster numbers and annotations of the heatmap refer to interaction affinities for interactions between clusters. Interaction terms range between 0 and 1, where 0 indicates a low likelihood of interaction and 1 indicates a high likelihood of interaction.

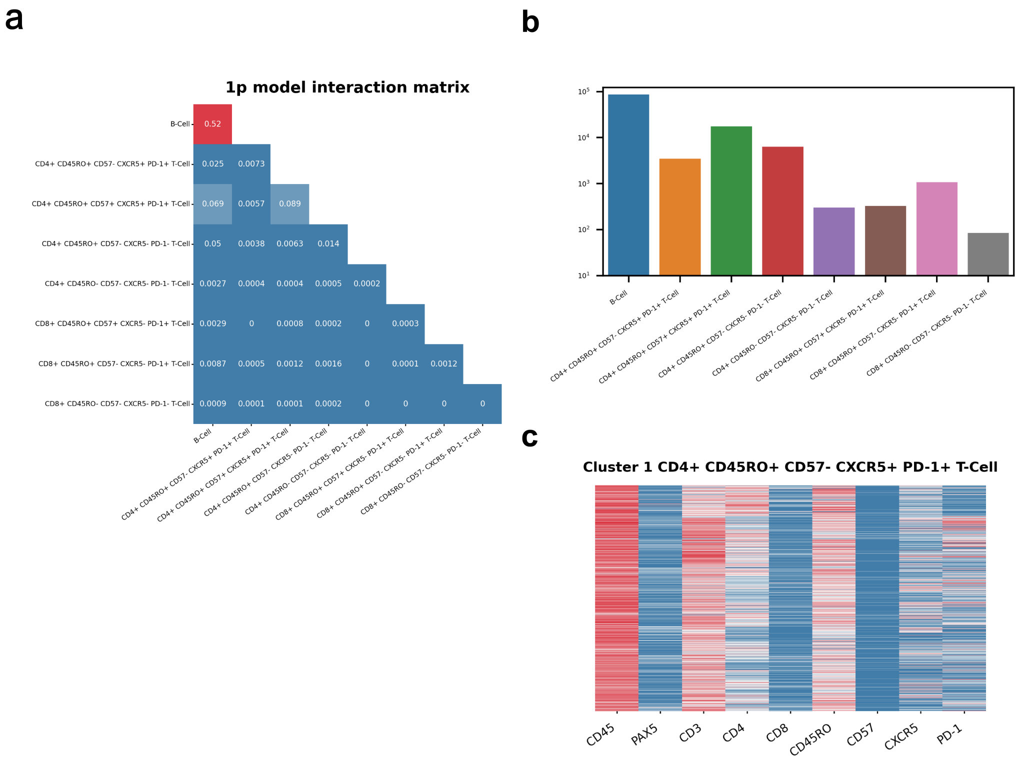

**Supplementary Figure 7**: Cell type annotation and cell-cell interaction analysis of DLBCL MIBI data using a 1p model. **a)** The interaction matrix for 29 patients with DLBCL with each cell of the matrix representing the probability distribution of an edge to be between two cell types in the HMRF. An edge represents cells of a cell type to be spatially proximal and interacting with cells of another cell type. **b)** The cell type distribution bar graph of the clustering results with log-scaled counts. **c)** An exemplar cluster heatmap of a CD4+ CD45RO+ CD57- CXCR5+ PD-1+ T cell from using the 1p model.

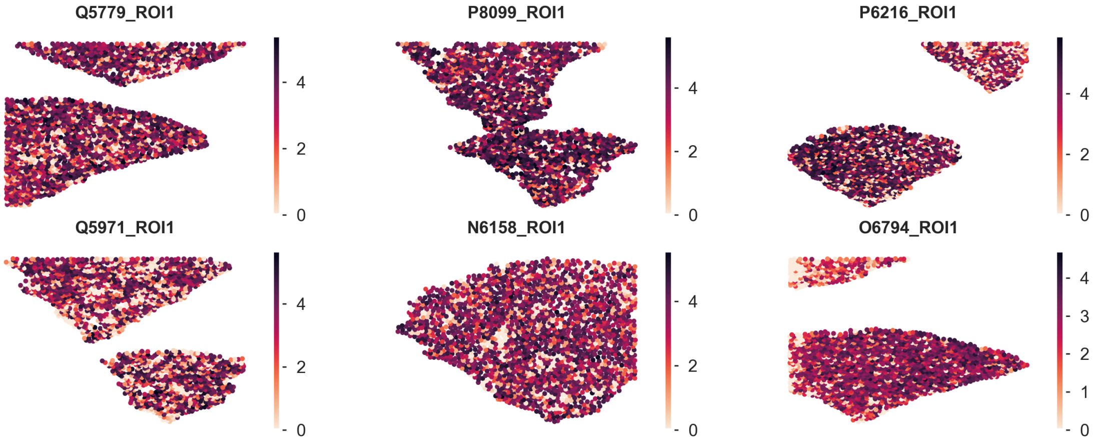

**Supplementary Figure 8**: Spatial distribution of the expression of B-cell lineage marker, PAX5, across cells in six patient samples. Color represents normalized intensity of expression.

#### 3 Supplementary Tables

**Supplementary Table 1**: Comparison of performance for forward simulated uniform and biased datasets. Means of v-measures are indicated in the table. See supplementary figures 2 and 3 for boxplot visualization.

**Supplementary Table 2**: Comparison of performance for spatial Gaussian mixture uniform and biased datasets. Means of v-measures are indicated in the table. See supplementary figures 4 and 5 for boxplot visualization.

**Supplementary Table 3**: Comparison of the performance of different clustering methods on semi-real biased and uniform datasets. Means of v-measures are indicated in the table. See figures 2 for boxplot visualization.

**Supplementary Table 4**: Comparison of the performance of different clustering methods on forward simulated biased and uniform datasets. The p-values are computed using the Friedman test. A p-value $< 0.01$ is considered significant.

**Supplementary Table 5**: Comparison of the performance of different clustering methods on spatial Gaussian mixture biased and uniform datasets. The p-values are computed using the Friedman test. A p-value $< 0.01$ is considered significant.

**Supplementary Table 6**: Comparison of the performance of different clustering methods on semi-real biased and uniform datasets. The p-values are computed using the Friedman test. A p-value $< 0.01$ is considered significant.

**Supplementary Table 7**: Comparison of the performance of different clustering methods on forward simulated biased and uniform datasets. The p-values are computed using the post-hoc nemenyi test. A p-value $< 0.01$ is considered significant.

**Supplementary Table 8**: Comparison of the performance of different clustering methods on spatial Gaussian mixture biased and uniform datasets. The p-values are computed using the post-hoc nemenyi test. A p-value $< 0.01$ is considered significant.

**Supplementary Table 9**: Comparison of the performance of different clustering methods on semi-real biased and uniform datasets. The p-values are computed using the post-hoc nemenyi test. A p-value $< 0.01$ is considered significant.
